# Flexibly-oriented double Cdc45-MCM-GINS intermediates during eukaryotic replicative helicase maturation

**DOI:** 10.1101/388470

**Authors:** Lu Liu, Yue Zhang, Jingjing Zhang, Jian-Hua Wang, Qinhong Cao, Zhen Li, Judith L. Campbell, Meng-Qiu Dong, Huiqiang Lou

## Abstract

The core of the eukaryotic helicase MCM is loaded as an inactive double hexamer (DH). How it is assembled into two active Cdc45-MCM-GINS (CMG) helicases remains elusive. Here, we report that at the onset of S phase, both Cdc45 and GINS are loaded as dimers onto MCM DH, resulting in formation of double CMG (d-CMG). As S phase proceeds, d-CMGs gradually mature into two single CMG-centered replisome progression complexes (RPCs). Mass spectra reveal that RPA and DNA Pol α/primase co-purify exclusively with RPCs, but not with d-CMGs. Consistently, d-CMGs are not able to catalyze either the unwinding or de novo DNA synthesis, while RPCs can do both. Using single-particle electron microscopy, we have obtained 2D class averages of d-CMGs. Compared to MCM DHs, they display heterogeneous, flexibly orientated and partially loosened conformations with changed interfaces. The dumbbell-shaped d-CMGs are mediated by Ctf4, while other types of d-CMGs are independent of Ctf4. These data suggest CMG dimers as bona fide intermediates during MCM maturation, providing an additional quality control for symmetric origin activation and bidirectional replication.

## Introduction

Eukaryotic cells exploit multilevel mechanisms to strictly control the initiation of DNA replication to achieve proper transmission of their genomes during cell proliferation. As an engine of the replication machinery for all eukaryotes, Mcm2-7 comprises the core of replicative helicase for unwinding the duplex genome (Bleichert *et al.*, 2017, Parker *et al.*, 2017). Intriguingly, Mcm2-7 (MCM) is loaded onto the double-stranded DNA (dsDNA) as a catalytically inactive, head-to-head double hexamer (DH) in G_1_ phase (Coster & Diffley, 2017, Evrin *et al.*, 2009, Li *et al.*, 2015, Remus *et al.*, 2009). Two co-activators, Cdc45 and the GINS heterotetramer (go ichi ni san, composed by Sld5, Psf1, Psf2 and Psf3), have been demonstrated to be essential for the assembly of holo-helicase CMG (Cdc45-MCM-GINS), which operates as a single 11-subunit complex moving along the leading strand during S phase (Costa *et al.*, 2011, Gambus *et al.*, 2006, Ilves *et al.*, 2010, Moyer *et al.*, 2006, O’Donnell & Li, 2018, Pacek *et al.*, 2006, Riera *et al.*, 2017, Yardimci *et al.*, 2010).

As cells proceed to S phase, the Dbf4-dependent Cdc7 protein kinase DDK phosphorylates the N-terminal tails of Mcm2/4/6 (Sheu & Stillman, 2006, Sheu & Stillman, 2010), triggering their interaction with Sld3-Cdc45 (Deegan *et al.*, 2016, Fang *et al.*, 2016, Heller *et al.*, 2011, Tanaka & Araki, 2013). This leads to the assembly of the Cdc45-MCM-Sld3 (CMS) platform. Then, Sld2 and Sld3 are phosphorylated by S-phase cyclin-dependent kinase (S-CDK), which promotes the formation and recruitment of the Sld2-Dpb11-Pol ε-GINS complex (Siddiqui *et al.*, 2013, Tanaka & Araki, 2013). It is conceivable that this step results in the replacement of Sld3 by GINS. These highly orchestrated events eventually produce the CMG complex, the core of RPCs (Abid Ali *et al.*, 2016, Bell & Labib, 2016, Bruck & Kaplan, 2015, Burgers & Kunkel, 2017, Sun *et al.*, 2016). Nevertheless, the details of how the MCM DH matures into two single CMG-centered RPCs (CMG/RPCs) remains unknown.

Previously, using a tandem affinity purification approach, we have purified the endogenous MCM DH from budding yeast (Quan *et al.*, 2015). In this study, through an expanded tandem affinity purification approach and glycerol sedimentation-velocity gradient centrifugation, we have identified various MCM-containing complexes formed as cells progress from G_1_ and then throughout the cell cycle. MCM persists in the dimeric form in the initial stage of Cdc45 and GINS association. Intriguingly, both Cdc45 and GINS exist in a dimerized form prior to being recruited onto the MCM DH on chromatin, leading to the assembly of a double CMG (d-CMG). With S phase progression, d-CMGs segregate gradually and this in turn leads to the appearance of single CMG/RPCs. The sequential changes of the components of various MCM complexes are revealed by mass spectrometry. The d-CMG fractions do not contain RPA and DNA Pol α/primase, which co-purify in single CMG (s-CMG)/RPCs exclusively. In contrast, both fractions have DNA Pol ε and Tof1/Mrc1/Csm3. Under the single-particle electron microscope (EM), our endogenous d-CMG fractions display a very different spectrum of conformations compared to the previously reported fly CMG complexes prepared by baculovirus mediated co-expression of recombinant Cdc45, four GINS and six MCM subunits (Costa *et al.*, 2014). These and other experiments reported here suggest that assembly and disengagement of double CMGs define a crucial step during helicase activation and replication initiation in vivo, as also recently reported with CMG assembled and activated in vitro using purified yeast proteins (Douglas *et al.*, 2018). Similarities and differences between our in vivo experimental findings and the yeast in vitro results will be discussed.

## Results

### MCM persists in the DH state upon the initial association of Cdc45 and GINS

Given that Cdc45 and GINS association is known to be capable of activating the MCM helicase (Ilves *et al.*, 2010), we first investigated the dimerization status of MCM upon the initial recruitment of Cdc45 and GINS in more detail. To this end, an extra copy of *MCM4* with a 5FLAG epitope was introduced into a yeast strain whose endogenous copy of *MCM4* was tagged with a calmodulin binding protein (CBP). This allowed isolation of a dimeric species of MCM through tandem affinity purification via calmodulin and anti-FLAG beads. The proteins eluted after each purification step were analyzed by western blotting. Psf2, a subunit of the GINS complex, coexisted with the MCM DH, as did Cdc45 (Figure 1A). Nonspecific association unlikely occurred under our tandem affinity procedure since no protein could be detected in the final eluates of the controls harboring only one of the epitope tags on *MCM4.* This result indicates that we have identified a native dimeric CMG complex in yeast cells as had been observed previously only in vitro (Costa *et al.*, 2014), and suggests that Cdc45 and GINS are recruited in the context of double hexameric MCM.

**Figure 1.**
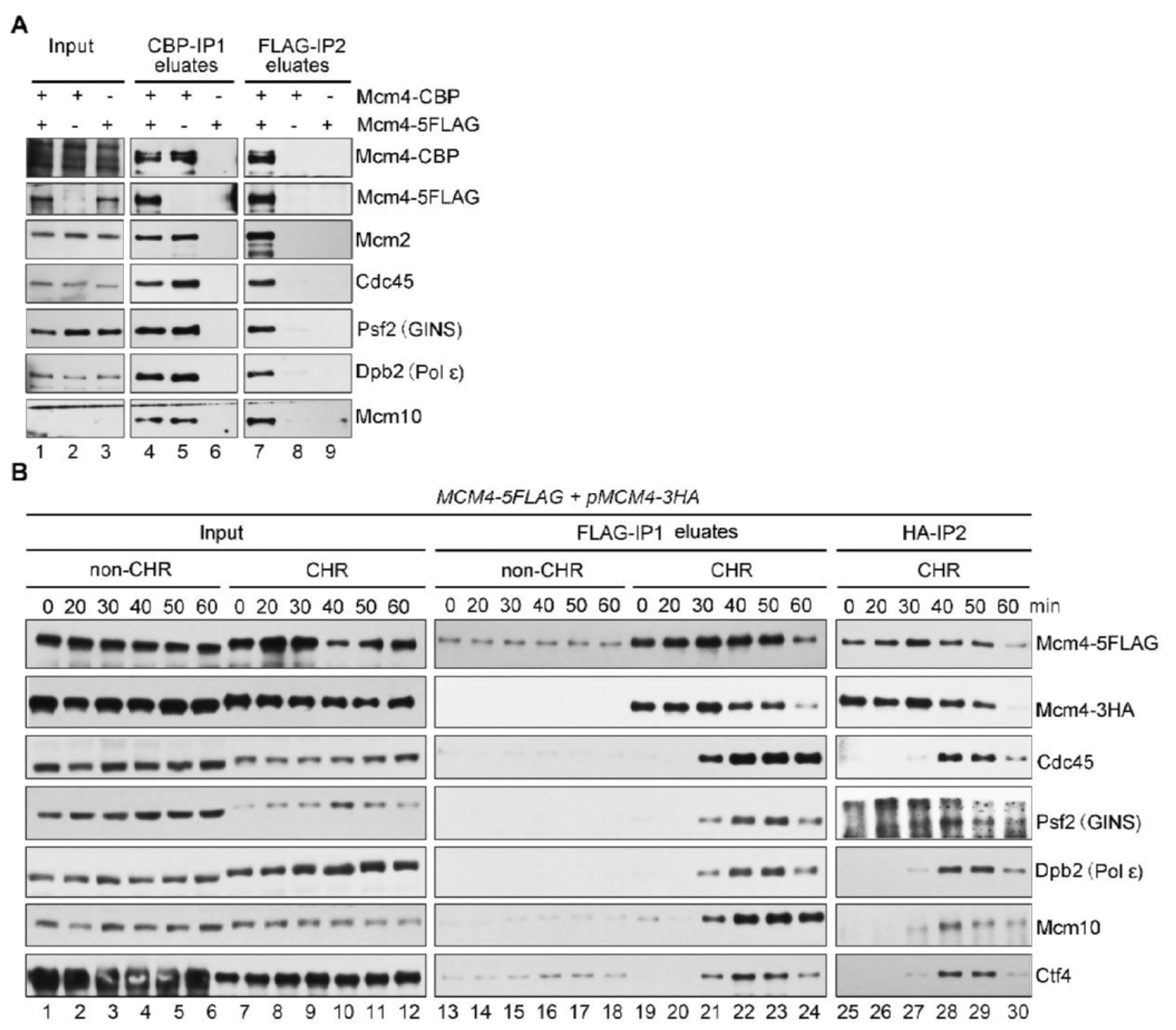
Identification of a double CMG complex during the S phase. (A) The *MCM4-CBP/pMCM4-5FLAG* cells (Strains LL94-1, Table S1) were cultured at 30°C and collected at OD_600_ about 1.0. Whole cell extracts (WCE) were prepared and subjected to tandem affinity purification via calmodulin and anti-FLAG M2 resins. After wash for at least three times, the bound fractions were eluted from beads by 3 mM EGTA (labeled as CBP-IP1 eluates) and 2 mg/ml FLAG peptides (labeled as FLAG-IP2 eluates), respectively. The eluted samples were resolved on an SDS-polyacrylamide gel and detected via immunoblots using the indicated antibodies. Strains (LL94-2 and LL6-1, Table S1) harboring a single tag (either CBP or 5FLAG) on Mcm4 were applied as controls. (B) The *MCM4-5FLAG/pMCM4-3HA* cells were grown, synchronized in G1 by α–factor (0 min) and released into S phase at 25°C for the indicated time. Spheroplasts were fractionated to the non-chromatin-bound (non-CHR) and chromatin-bound (CHR) protein fractions. Mcm4-5FLAG and then Mcm4-3HA were precipitated consecutively in a similar procedure mentioned above. After wash for three times, the proteins specifically associated with beads were eluted by 2 mg/ml of FLAG peptide or boiled directly (for HA-IP) before western blotting.

### Assembly and segregation of dimeric CMGs during S phase

To further confirm the formation of dimeric CMGs, we prepared chromatin-bound (CHR) and non-chromatin-bound (non-CHR) fractions from cells synchronized in G1 (0 min) or released into S phase for 20, 30, 40, 50 or 60 min. To rule out possible artifacts associated with the pair of tags used in Figure 1A, the dimeric form of MCM was obtained by using a second set of affinity tags (FLAG and HA). MCM DHs, i.e., double labeled FLAG/HA Mcm2-7 complexes, were detected exclusively in the chromatin fraction (Figure 1B). The MCM DH already appeared in G1 phase, before release into S phase. However, no additional proteins were detected in the complexes in G_1_. After release into S phase, other initiation factors including Dpb2 (a subunit of DNA Pol ε), Cdc45 and GINS were first detected in the chromatin-associated MCM complex after about 30 min release. The amounts of these initiation factors peaked at ~40 min and was coincident with the decline in the level of the MCM DH (Figure 1B). These results show that there is a bona fide dimeric CMG status before gradual dissolution during S phase progression.

To further validate and characterize the different species of the MCM-containing complexes during the cell cycle, we next subjected the FLAG peptide eluates from the first immunoprecipitation (i.e. FLAG-IP) of the CHR fraction mentioned above on a 10-30% glycerol sedimentation/velocity gradient. In G_1_ phase, only the MCM DH, peaking at fractions 15-17, was detected (Figure 2A). This fraction sedimented more rapidly than a 669 kDa protein standard (fraction 13), identifying it as a double hexameric MCM (theoretically 1211 kDa), as shown previously (Quan *et al.*, 2015). When cells entered S phase after 30 min, the MCM-containing complexes appeared to co-sediment with Cdc45 and GINS over a broader range. The peak of Cdc45 (fraction 15) coincided with the peak of MCMs in all time points. Although the separation is not complete, it seems that there are two distinct populations of complexes, one migrating in the lower part of the gradient relative to MCM DH (Figure 2A, fractions 10-13) and the other migrating at higher positions than MCM DH (fractions 18-21). Notably, Mcm4-3HA tended to co-sediment with Mcm4-FLAG in the higher gradient fractions, suggesting that this portion likely represents the dimeric CMG species detected in Figure 1. In agreement with this, Mcm10, an essential initiation factor known to preferentially bind the MCM DH (Douglas & Diffley, 2016, Quan *et al.*, 2015), primarily enriched in the higher density gradients (fractions 18-21) as well. These results imply that the fast-sedimenting MCM complex may be the dimeric species of CMG.

**Figure 2.**
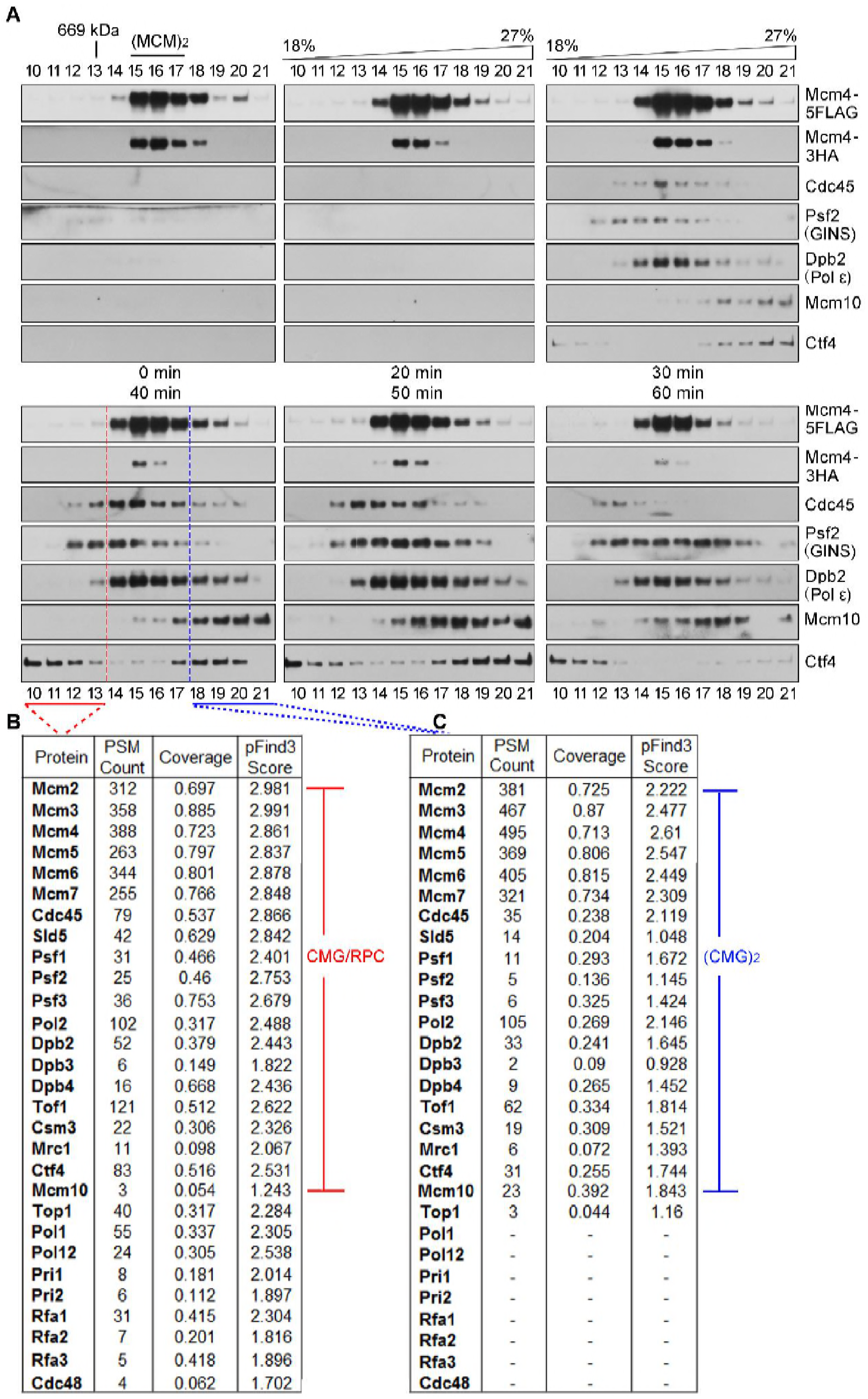
Dynamic changes of the MCM double hexamer throughout the cell cycle. (A) The *MCM4-5FLAG/pMCM4-3HA* cells were synchronized and collected as described in Fig. 1B. The disparate forms of the MCM complexes were isolated from CHR fractions via one step purification (i.e., FLAG-IP and FLAG peptide elution) followed by 10-30% glycerol density gradient centrifugation. After centrifugation at 79,000g for 16h in Hitachi CP100NX with a P55ST2 rotor, total 4.8 ml sample was equally divided into 24 fractions (1-24, from top to bottom). 25 μl of each fraction was analyzed by immunoblotting. The fraction number was indicated for each lane. (MCM)_2_ and (CMG)_2_ represent the dimeric forms of MCM and CMG, respectively. (B) Mass spectra of the slow- and fast-sedimenting fractions. Fractions 10-13 and 18-21 were pooled before precipitating the proteins for LC-MS/MS analysis. The total number of identified peptides, coverage and pFind3 score are summarized.

To further test this possibility, we then determined the composition of these CMG complexes by mass spectrometry. The slow-sedimenting fractions (10-13) and fast-sedimenting fractions (18-21) were pooled separately prior to trypsin digestion. Besides the essential initiation factors (Yeeles *et al.*, 2015), other replication progression factors including the fork protection complex (Tof1-Mrc1-Csm3) required for efficient DNA replication (Yeeles *et al.*, 2015), were also detected in both S-phase-specific MCM complexes. Strikingly, RPA (Rfa1-Rfa2-Rfa3), DNA Pol α and primase (Pri1 and Pri2) presented only in the slow-sedimenting complex (Figure 2B), not in the fast-sedimenting one (Figure 2C). Given that the loading of RPA and Pol α/primase requires single-stranded DNA, these results implicate that the slow-sedimenting and fast-sedimenting species of the S-phase-specific MCM complexes might represent the active s-CMGs and inactive d-CMGs, respectively. Moreover, the components identified in the slow-sedimenting complex correlate well with previous systematic mass spectra of RPC and its associated factors (Gambus *et al.*, 2006). Taken together, these data suggest that the MCM DH is initially assembled into a dimeric form of CMG before transition into two monomeric active CMGs associated with additional fork progression proteins.

### Cdc45 and GINS are loaded in a dimerized form

Next, we asked how double CMGs are assembled in yeast cells. Given the fact that each active CMG contains one Cdc45 and GINS, we speculated that there should be two molecules of Cdc45 and GINS in a double CMG. To understand the mode of their recruitment, we constructed a strain containing two copies of Cdc45 tagged with a 5FLAG and a 13MYC, respectively. First, Cdc45-5FLAG was precipitated from whole cell extracts. Cdc45-13MYC was clearly detected in the precipitates, but not in the mock IPs (Figure 3A). To examine whether intermolecular interaction of Cdc45 occurs in the context of chromatin, we next repeated FLAG-IPs in both non-CHR and CHR fractions. Interestingly, Cdc45-13MYC co-precipitated with Cdc45-5FLAG in both cases (Figure 3B). We further analyzed the Cdc45 complexes eluted from FLAG-IPs by glycerol gradient centrifugation. In the non-CHR fraction, Cdc45 sedimented very slowly and peaked at the same fractions as aldolase (158 kDa), close to the predicted molecular weight of a Cdc45 dimer (148 kDa). This indicates that Cdc45 very likely exists as a dimer prior to chromatin association. Meanwhile, in the CHR fraction, Cdc45-5FLAG co-sedimented with Cdc45-13MYC, MCM, GINS and Dpb2 to a similar range of density gradients as putative double CMGs shown in Figure 2A (Figure 3C). Because the chromatin-bound (CHR) fraction was released as a complex via benzonase, the isolated complexes represent protein-protein interactions and not just indirect association through DNA.

**Figure 3.**
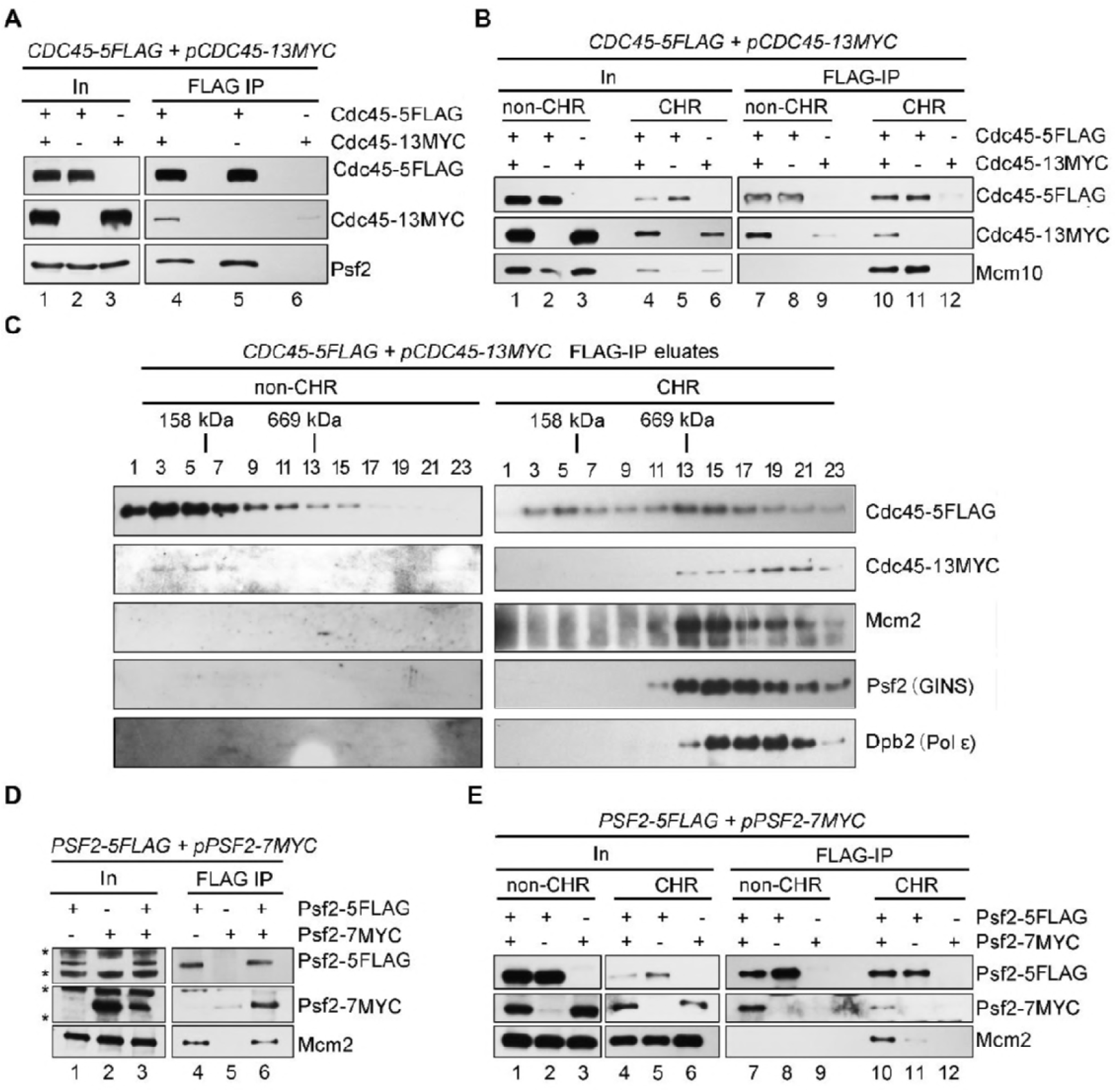
Cdc45 and GINS are loaded onto the MCM double hexamer as dimers. (A, B) Cdc45-5FLAG was precipitated via M2 beads from WCE (A), non-CHR or CHR (B) fractions of the *CDC45-5FLAG/pCDC45-13MYC* cells (Strain LL85-1, Table S1). Co-precipitated proteins were detected via immunoblots against the indicated antibodies. (C) Glycerol density gradient separation of the Cdc45-containing complexes. The *CDC45-5FLAG/pCDC45-13MYC* cells were cultured at 30°C and released into S phase for 40 min at 25°C after α–factor synchronization. Cells were then collected and fractionated. The Cdc45-containing complexes were purified by subjecting the Cdc45-FLAG eluates of non-CHR and CHR fractions onto a 10-30% glycerol density gradient as described in Figure 2. (D, E) Pfs2-5FLAG was precipitated via M2 beads from WCE (D), non-CHR or CHR (E) fractions of the *PSF2-5FLAG/pPSF2-7MYC* cells (LL67-1, Table S1). The precipitates were subjected to immunoblotting. Cross bands are labeled by asterisks.

Using a similar strategy, we were able to show that Psf2 also has intermolecular interaction (Figures 3D and 3E) and exists as a dimer before being loaded onto chromatin as well (Figure 4A). In contrast, MCM presents as a single hexamer before being loaded onto chromatin. It is also worth noting that Ctf4 co-purified with GINS, in agreement with the previous report that Ctf4 binds GINS directly (Gambus *et al.*, 2009). Given the fact that Ctf4 is a trimeric hub (Simon *et al.*, 2014, Villa *et al.*, 2016), the dimerization of GINS could be mediated by Ctf4. To test this possibility, we examined the oligomeric status of GINS in the *ctf4Δ* cells. The sedimentation of GINS in both non-CHR and CHR bound fractions was unchanged in the absence of Ctf4 (Figure 4B, compare to 4A). This result indicates that GINS and CMG dimers are not formed by Ctf4 (e.g., artificially during the purification). Taken together, these data suggest that both Cdc45 and GINS are recruited onto the MCM DH as dimers, which results in the initial assembly of d-CMGs on chromatin.

**Figure 4.**
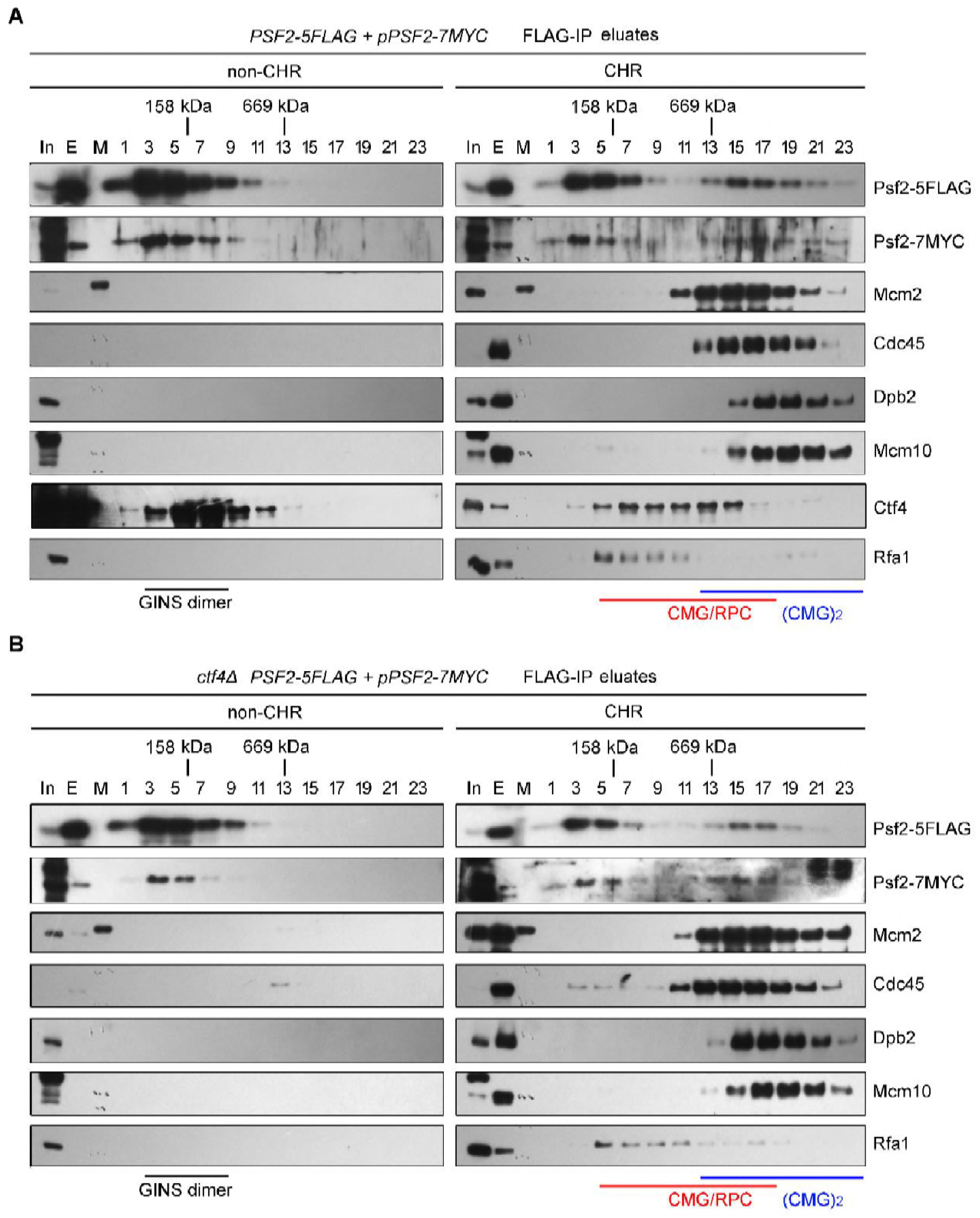
Both GINS and CMG dimers are independent of the Ctf4 trimer. (A) GINS are loaded onto chromatin in a dimerization form. The *PSF2-5FLAG/pPSF2-7MYC* cells in WT background were synchronized and collected after 40 min release at 25°C as in Figure 2. The Psf2-5FLAG complexes were precipitated from non-CHR or CHR fractions and the eluates were then subjected to glycerol density gradient separation. Each density fraction was analyzed via immunoblots against the indicated antibodies. (B) Dimerization of both GINS and CMG does not depend on Ctf4. The Psf2-5FLAG complexes from a *ctf4Δ* background were isolated and analyzed as described above.

### D-CMGs have no helicase and replication activities

Above results imply that d-CMG may represent an intermediate status between the MCM DH and s-CMG. To test this hypothesis, we next measured DNA helicase (Figure 5A) and de novo synthesis activities (Figure 5B) in each fraction of density gradient centrifugation. Fractions 11-17 displayed clear unwinding activity on a 5’-^32^P-labeled partial duplex DNA (Y-form DNA) in the presence of ATP at 30°C (Figure 5A). The substrates disappeared in fractions 7-11 probably due to degradation by nucleases, which are often associated with replisome. Then, the unlabeled version of the same Y-form DNA substrate was used as a template to examine the in vitro DNA synthesis activity. The products of replicated DNA were monitored by the incorporation of α-^32^P-dATP through autoradiography after separation on a denatured polyacrylamide gel. As shown in Figure 5B, in the presence of all four NTPs and dNTPs, fractions 11-17 were also able to catalyze the synthesis of the full-length (85-mer) DNA, indicating an efficient synthesis activity. Both helicase and replication activities peaked around fraction 15. It is worth emphasizing that no primers were included in the reactions and the RNA-dependent extension of DNA Pol α is usually limited to 10-12 nucleotides (Perera *et al.*, 2013). Therefore, the appearance of 85-mer products containing α-^32^P-dAMP should reflect at least three kinds of essential activities including helicase, primase and polymerase in the DNA replication process. These results are consistent with the presence of CMG, Pri1/2, DNA Pol α, and Pol ε in these putative RPC fractions as revealed in mass spectra (Figure 2B). To exclude the possibility that α-^32^P-dAMP is incorporated by contaminating terminal deoxynucleotidyl transferase (TdT) activity, we incubated TdT with the unlabeled Y-shaped substrate in the presence of α-^32^P-dATP. Products much longer than 85-mer were detected (Figure 5C, lane 6), which were very sensitive to single-stranded DNA specific S1 nuclease (lanes 7 and 8). However, no products longer than 85-mer were observed for the putative RPC fractions (Figure 5C, lane 4). More importantly, 85-mer products can only be digested if they are boiled prior to S1 treatment (Figure 5C, compare lanes 2 and 5). These results allow us to conclude that the products replicated by the RPC fractions (fractions 11-17, Figure 5B) are duplex DNA. In stark contrast, there were neither unwound (Figure 5A) nor replicated DNA products (Figure 5B) detectable in the fast-sedimenting fractions (19-23). Taken together, these results argue that the slow-sedimenting complexes are single active CMG/RPCs, while the fast-sedimenting complexes may represent the immature d-CMGs.

**Figure 5.**
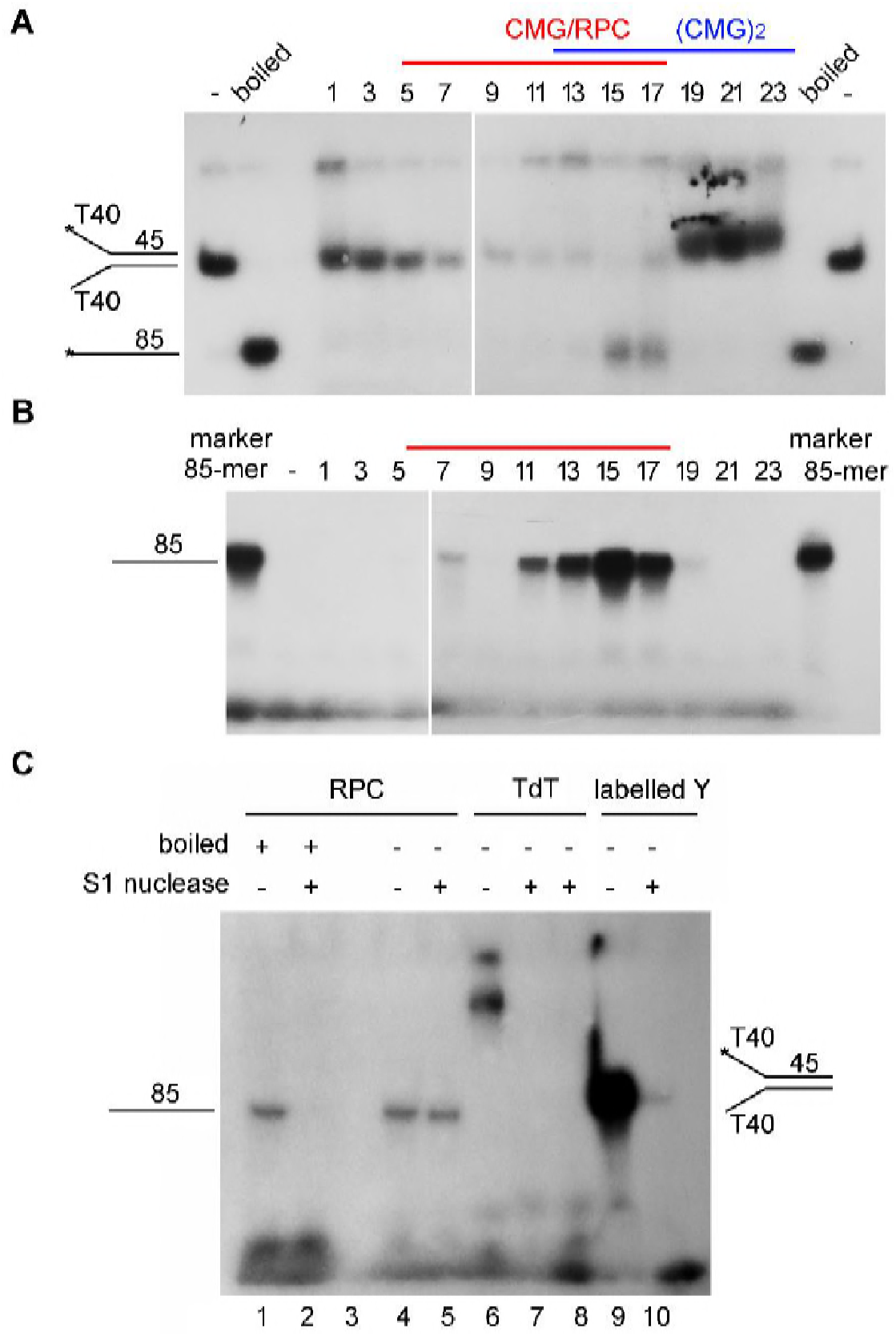
The fast-sedimenting fractions have few helicase and synthesis activities. (A) In vitro helicase assay. The Psf2-5FLAG complexes were purified exactly as described in Figure 4. Each fraction from glycerol gradient centrifugation was subjected to in vitro helicase assays as described in Experimental Procedures. A Y-shaped duplex DNA labeled at 5’-end with ^32^P was purified and used as a substrate. The products were analyzed by a native 8% polyacrylamide gel followed by autoradiography. Boiled substrates were loaded to indicate the migration of an 85-mer oligonucleotide. (B) In vitro DNA synthesis assay. Each fraction was also applied to the same Y-form substrate without ^32^P-labelling for measuring DNA synthesis activity in the presence of all four kinds of NTPs and dNTPs including α-^32^P-dATP at 30°C for 60 min. The reactions were quenched and resolved by a 20% polyacrylamide gel containing 8 M urea. The synthesized products were detected by incorporation of ^32^P-dAMP in autoradiography. A ^32^P-labeled 85-mer was loaded as a size marker. (C) The ^32^P-dAMP incorporated products by RPC are resistant to S1 nuclease. In vitro DNA synthesis assays were performed as described above for both RPC fractions (11-17) and terminal deoxynucleotidyl transferase (TdT) enzymes. The final products were treated by S1 nuclease with or without boiling. The pre-labelled Y-DNA was digested by S1 nuclease as a control.

### D-CMGs display heterogeneous and rotated conformations

To directly observe the dimeric form of CMG, we then examined the CMG complexes from the fractions of gradient centrifugation using a transmission electron microscope after negative staining with uranyl acetate. The majority of the CMG particles were homogeneous in size (20-23 nm) with a noticeable central channel from the top/bottom view (Figures 6A and 6B), in good agreement with the high resolution structure of s-CMGs as reported very recently (Figures 6C) (Georgescu *et al.*, 2017, Sun *et al.*, 2016, Yuan *et al.*, 2016). Interestingly, DNA Pol ε, Ctf4 and other components co-purified with s-CMGs (Figure 6B), representing relatively stable parts of RPCs. Consistent with recently resolved EM structures (Sun *et al.*, 2015, Zhou *et al.*, 2017), Pol ε associated with CMG through the C-terminal tier of the MCM complex and Ctf4 associated through GINS (Figure 6D). These results corroborate that we have successfully purified the endogenous CMG complexes from yeast cells using our tandem affinity approach. In addition to s-CMGs, a proportion of particles appeared to have a markedly larger size (~35 nm), approximately twice the size of s-CMGs (Figure 6A). Unlike the MCM DHs and s-CMGs, the putative d-CMGs display markedly heterogeneous conformations, suggesting increased flexibility (green squares, Figure 6B). This is in contrast to the d-CMG reconstituted in vitro from the purified fruit fly proteins associates stably with each other through the MCM N-termini just as in its precursor MCM DH (Costa *et al.*, 2014). Moreover, the class averages of our representative d-CMG species showed that the two component CMGs are positioned in several different orientations (Figures 6B and 6E). A sub-population of these d-CMGs, which we refer to as “dumbbell-shaped”, revealed two MCM hexamers that appear to have detached from each other. Their association could be mediated by other components such as Ctf4 (Figures 6E).

**Figure 6.**
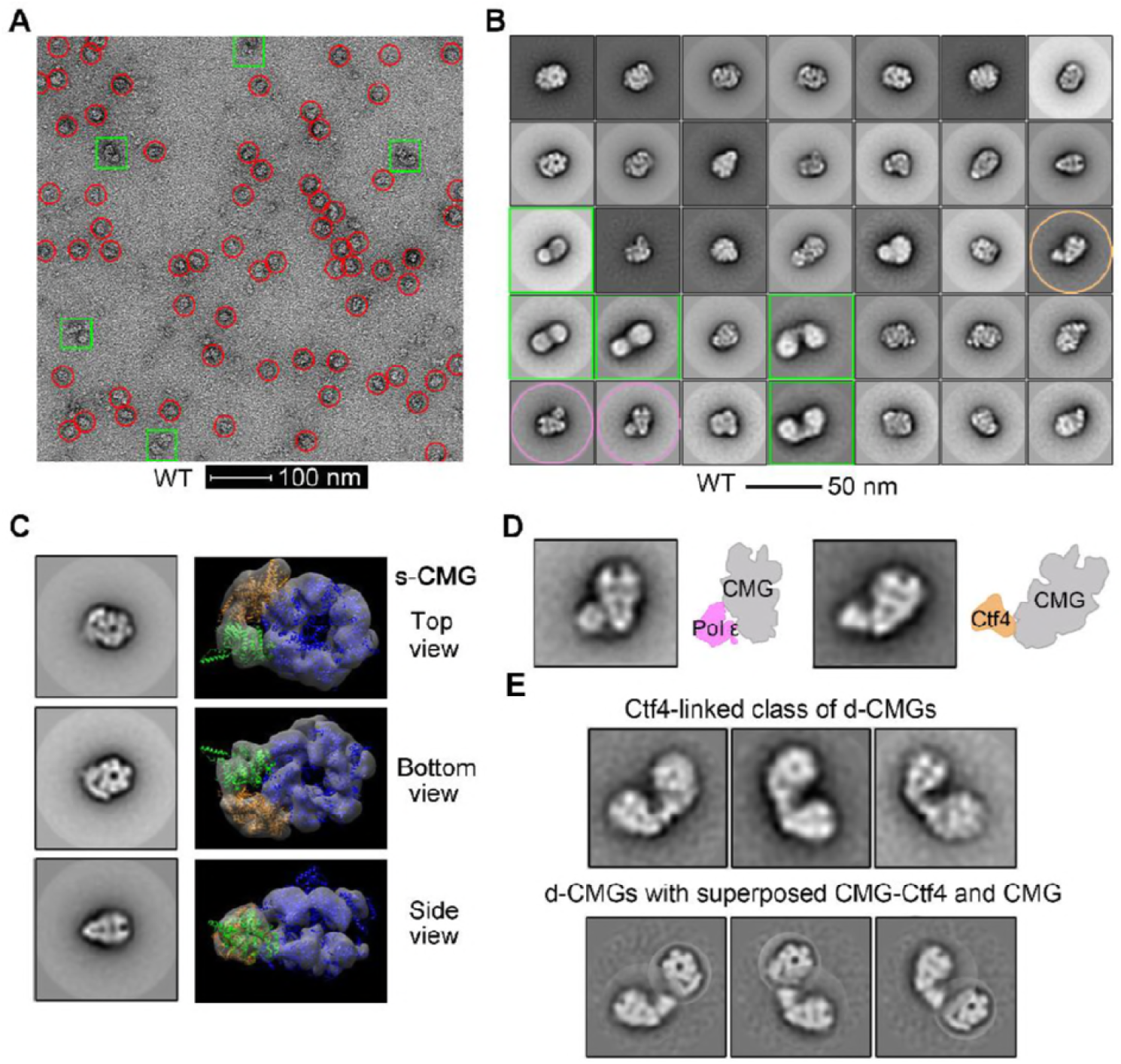
Single-particle EM analysis of the negatively stained CMG complexes. (A) A representative electron micrograph of the endogenous CMG complexes isolated from the *CDC45-5FLAG* cells (LL85, Table S1) through the same purification procedure as described in Figure 4. The single (s-CMG) and double (d-CMG) CMG particles are highlighted by red circles and green squares, respectively. (B) 2D class averages of all types of CMG particles (38,787 in total). (C) S-CMG particles with top/bottom and side views. (D) S-CMG particles containing DNA Pol ε or Ctf4. (E) The dumbbell-shaped d-CMG particles (824 among total 6,445 d-CMGs) with superposed CMG-Ctf4 and CMG.

### Ctf4-independent types of d-CMG

Given that Ctf4 is a trimeric hub directly associating with GINS, to exclude the artifactual formation of oligomeric CMGs during purification, we next monitored d-CMG species isolated in the *ctf4Δ* background. Indeed, deletion of Ctf4 abolished the “dumbbell-shaped” d-CMGs (Figures 7A), indicating that this type of d-CMGs is loosely connected by Ctf4. However, as shown in Figure 7A, in the absence of Ctf4, many other types of d-CMGs persisted, consistent with the observations in Figure 4B. These indicate that d-CMGs are bona fide supercomplexes coexisting with s-CMGs.

**Figure 7.**
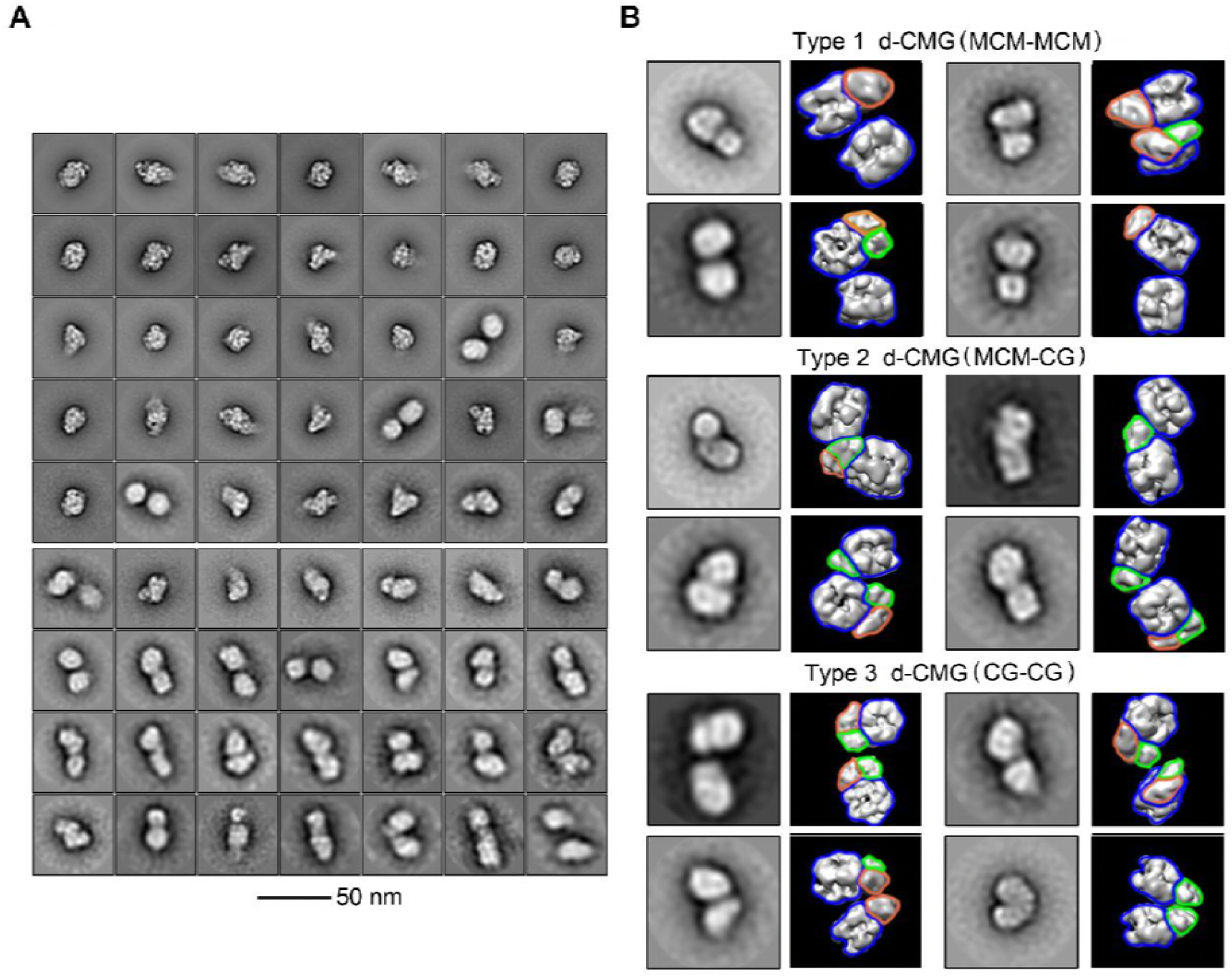
Ctf4-independent d-CMG species. (A) 2D class averages of the CMG particles (43,820 in total) purified endogenously from the *ctf4Δ* cells (LL-163, Table S1). (B) D-CMG particles (4,389 in total) with model docking of two s-CMG structures (PDB code 3JC5). The putative interfaces of different types of d-CMGs are indicated in parenthesis. Blue, MCM; Green, Cdc45; and Orange, GINS. S-CMGs fit well with the density map. Due to the orientation variation in the d-CMG complexes, the density for them is often fragmented. In addition, there appears to be extra density that could be attributed to other replication factors.

A preliminary 2D average nicely resolved densities for s-CMGs, whereas the CMGs from the d-CMG particles from the *ctf4Δ* cells were mostly smeared out (Figures 7A). A further examination indicates that there are multiple types of d-CMGs with different interfaces mediated via MCM, Cdc45 or GINS for instance (Figure 7B). These results imply that in our purified endogenous d-CMGs, the tight association between the two single MCM hexamers (Evrin *et al.*, 2009, Li *et al.*, 2015, Remus *et al.*, 2009) might have undergone conformational changes/rotations, resulting in partial disruption of the tightly associated MCM-MCM within the DH, in agreement with the very recent observation in vitro (Douglas *et al.*, 2018). Taken together, these data suggest that d-CMGs likely undergo multiple conformational changes accompanying the cell-cycle-regulated association of dimeric Cdc45, GINS and/or other firing factors on the path to maturation into single active CMG helicases.

## Discussion

Since the discovery of the MCM DH assembly during the licensing stage, how two single hexamers at an origin are simultaneously activated to achieve bidirectional DNA replication becomes a key conundrum in eukaryotic DNA replication field. Here, we provide in vivo evidence to support a bona fide dimeric CMG intermediate in yeast cells with some unanticipated characteristics, which may provide important insight to bidirectional replication and helicase remodeling.

Cdc45 and GINS have been well established as essential co-activators for the core MCM hexamer. Therefore, the finding that both Cdc45 and GINS are recruited to the MCM DH as dimers provides an additional mechanism, likely instrumental to achieving simultaneous activation of both MCM hexamers and bidirectional DNA replication from each origin. In the MCM DH, the two hexamers associate head to head with abutting N terminal tiers. Interestingly, single CMG translocates with the MCM N-tier ahead of the C-tier (Douglas *et al.*, 2018, Georgescu *et al.*, 2017). Based on this important finding, it has been proposed that the two single CMGs must pass each other on opposite strands during initiation, providing an elegant fail-safe mechanism to ensure complete bidirectional replication of origin DNA. Our data supporting the assembly of a dimeric CMG by both Cdc45 and GINS dimers provides an additional layer of quality-control at an even earlier stage (i.e., pre-initiation stage).

Cdc45 from other organisms has been observed to be able to form dimers in vitro (Chang *et al.*, 2007, Kamada *et al.*, 2007). Interestingly, Sld3, the hub mediating CMG assembly, can be dimerized through its chaperone Sld7 in an antiparallel manner in vitro (Itou *et al.*, 2015). Moreover, two copies of recombinant archaeal GINS and Cdc45 may form a stable complex (Xu *et al.*, 2016). DNA Pol ε, forming a CDK-dependent pre-loading complex with GINS (Muramatsu *et al.*, 2010), may be integrated as a dimer mediated by Dbp2 as well (Dua *et al.*, 2000, Sengupta *et al.*, 2013). All these in vitro observations, together with our finding that yeast Cdc45 and GINS exist in dimers in vivo, arguing for an evolutionarily conserved symmetric activation of the two MCM hexamers on an MCM DH (Swuec & Costa, 2017, Watson & Crick, 1953).

The endogenous d-CMGs identified in this study exhibit heterogeneous and flexible conformations, which is distinct from the d-CMG/DNA complexes prepared by reconstitution of 11 CMG baculovirus expressed CMG subunits and DNA reported previously (Costa *et al.*, 2014). The in vitro reconstituted *Drosophila melanogaster* dimeric CMG is homogenously oriented head-to-head through tight association between MCM N-termini as in the MCM DH. We propose that these observed conformations could represent different stages of d-CMG. Supporting this, only a small proportion of the CMG particles exists as dimers in both studies. It is also not surprising that double CMG is a flexible and transient intermediate given the starkly different structures of its precursor MCM DH and its product s-CMG observed to date. Therefore, the dimeric CMG complexes captured in the in vitro reconstitution might represent an initial state, whereas our d-CMGs represent later stages during remodeling. It will be interesting to find out the exact underlying reasons for such differences in the future.

According to the high resolution CMG structure obtained recently, Cdc45 and GINS finally position near the Mcm2-Mcm5 gate, which orients near oppositely within the MCM DH (Georgescu *et al.*, 2017, Sun *et al.*, 2016, Yuan *et al.*, 2016). Therefore, it is conceivable that dimerized Cdc45 and GINS could help to induce conformational changes (e.g., axial rotation) of the two MCM rings, thereby weakening or interrupting the tight head-to-head association within a MCM DH as observed by Diffley’s group in vitro (Douglas *et al.*, 2018). Such a weakened MCM-MCM association may be difficult to detect at the CMG stages in some certain conditions despite similar strategies are used (Miyazawa-Onami *et al.*, 2017). In accordance with this, we found that two MCM rings have detached and positioned in different orientations in most types of d-CMGs. It indicates that the two tilted and twisted MCM hexamers have undergone rotation (Li *et al.*, 2015). Speculatively, the relative movements of the two MCM single hexamers could simultaneously induce the melting of the duplex DNA embraced inside the MCM DH. All these possibilities are worthy to be further tested in future.

### Experimental procedures

#### Strain and plasmid construction

*Saccharomyces cerevisiae* strains and plasmids used in this study are listed in Table S1 and S2, respectively.

Cell synchronization, whole cell extract preparation and chromatin fractionation, immunoprecipitation (IP) were performed as previously described (Quan *et al.*, 2015).

#### Glycerol density gradient centrifugation

The native protein complexes in the peptide eluates after FLAG-IPs were concentrated and applied to the top of a 10-30% glycerol gradient in elution buffer without protease inhibitors. The gradients were centrifuged in a P55ST2 swinging bucket rotor (Hitachi ultracentrifuge) at 79,000g for 16 h using slow deceleration. Following centrifugation, 24 fractions (200 μl each) were collected manually from top to bottom of the gradient. As molecular weight markers, a mixture of bovine serum albumin (68 kDa), aldolase (158 kDa) and thyroglobulin (669 kDa) was centrifuged in a separate tube. The fractions containing different species of the MCM complexes were pooled and processed for mass spectrometry, in vitro helicase/replication and single-particle EM analysis described below.

#### Helicase assays

The helicase activity was measured using a 5’-^32^P-labeled 85 bp duplex DNA substrate bearing a single-stranded 3’-dT(_40_) tail with some modifications from (Xia *et al.*, 2015). Briefly, each reaction (37 μl) contains 0.5 nM 5’-^32^P-labeled Y-shaped DNA and 30 μl protein fraction collected from glycerol gradient centrifugation in a final helicase buffer (25 mM HEPES-KOH (pH 7.6); 150 mM potassium glutamate; 10 mM magnesium acetate; 0.1 mM EDTA; 2 mM DTT; 2 mM ATP). Reactions were conducted at 30°C for 60 min before addition of 4 μl quench buffer (200 mM EDTA, 1% SDS and 0.1% bromophenol blue). Products were then separated on a native 8% polyacrylamide gel in 0.5 × TBE before autoradiography.

#### De novo DNA synthesis and S1 nuclease-resistant assays

The DNA synthesis activity of each fraction from glycerol gradient centrifugation was measured using an unlabeled version of the Y-shaped DNA used in the helicase assays. Synthesis reactions (40 μl each) contain 0.5 nM unlabeled Y-form DNA and 33 μl of each fraction from glycerol gradient centrifugation in a final synthesis buffer (40 mM HEPES-KOH (pH 7.6); 150 mM potassium glutamate; 10 mM magnesium acetate; 2 mM DTT; 2 mM ATP) plus four NTPs (200 μM each), four dNTPs (40 μM dGTP/dCTP/dTTP and 4μM dATP) and 40 nM α-^32^P-dATP. Reactions were conducted at 30°C for 60 min.

For terminal deoxynucleotidyl transferase (TdT) assay, the reactions (30 μl each) contain 0.5 nM unlabeled Y-form DNA and 0.17 U/μl TdT (New England Biolabs) in a final buffer with 1×TdT reaction buffer, 1 μM dATP and 55 nM α-^32^P-dATP. Reactions were conducted at 37°C for 60 min before being inactived at 75 °C for 20 min.

For S1 nuclease treatment, the synthesized products by the RPC fractions or TdT were subjected to S1 nuclease digestion before analysis. S1 nuclease (final concentration 1 U/μl) was incubated at 25°C for 30 min with 50 μl synthesis reaction with or without prior boiling treatment. The reactions were stopped by addition of 6 μl quench buffer (200 mM EDTA and 0.1% bromophenol blue). All reaction products were separated on a 20% polyacrylamide gel containing 8 M urea in 1 × TBE before autoradiography.

#### MS sample preparation

Proteins were precipitated with 25% trichloroacetic acid (TCA) for at least 30 minutes on ice. The protein pellets were washed twice with 500 μl ice-cold acetone, air dried, and then resuspended in 8 M urea, 20 mM methylamine, 100 mM Tris, pH 8.5. After reduction (5 mM TCEP, room temperature, 20 min) and alkylation (10 mM iodoacetamide, room temperature, 15 min in the dark), the samples were diluted to 2 M urea with 100 mM Tris, pH 8.5 and digested with trypsin at 1/50 (w/w) enzyme/substrate ratio at 37°C for 16-18 hr. The digestion was then stopped by addition of formic acid to 5% (final concentration).

#### LC-MS/MS analysis

All protein samples were analyzed using an EASY-nLC 1000 system (Thermo Fisher Scientific, Waltham, MA) interfaced with a Q-Exactive mass spectrometer (Thermo Fisher Scientific). Peptides were loaded on a pre-column (75 μm ID, 4 cm long, packed with ODS-AQ 12 nm-10 μm beads) and separated on an analytical column (75 μm ID, 12 cm long, packed with Luna C18 1.9 μm 100 Å resin) with a 60 min linear gradient at a flow rate of 200 nl/min as follows: 0–5% B in 2 min, 5–30% B in 43 min, 30–80% B in 5 min, 80% B for 10 min (A = 0.1% FA, B = 100% ACN, 0.1% FA). Spectra were acquired in data-dependent mode: the top ten most intense precursor ions from each full scan (resolution 70,000) were isolated for HCD MS2 (resolution 17,500; NCE 27) with a dynamic exclusion time of 30 s. The AGC targets for the MS1 and MS2 scans were 3e6 and 1e5, respectively, and the maximum injection times for MS1 and MS2 were both 60 ms. Precursors with 1+, more than 7+ or unassigned charge states were excluded.

The MS data were searched against a Uniprot *S. cerevisiae* protein database (downloaded from Uniprot on 2013-04-03) using an updated version of pFind (Chi *et al.*, 2015) with the following parameters: instrument, HCD-FTMS; precursor mass tolerance, 20 ppm; fragment mass tolerance 20 ppm; open search mode; peptide length, minimum 6 amino acids and maximum 100 amino acids; peptide mass, minimum 600 and maximum 10,000 Da; enzyme, Trypsin, with up to three missed cleavage sites. The results were filtered by requiring FDR<1% at the spectral level and spectra count ≥ 2.

#### Electron microscopy

The CMG complexes were isolated from the peak fractions from glycerol density gradient centrifugation and concentrated by ultrafiltration. Negative staining of the samples deposited on carbon-coated grids was conducted with 2% uranyl acetate. Grids were examined using an FEI Tecnai F20 microscope operated at 200 kV, and images were recorded at a nominal magnification of 50,000 × using a 4k × 4k charge-coupled device (CCD) camera (UltraScan 4000, Gatan), resulting in a 1.7 Å pixel size at the specimen level.

#### Image processing and atomic docking

EMAN2 was used for manual particle-picking and micrograph-screening (Tang *et al.*, 2007). The 2D classification, 3D classification and 3D refinement were performed using RELION1.4 (Scheres, 2012). Artificial CMG dimers were generated by relating the two CMG atomic models (PDB code: 3JC5) in UCSF Chimera (Pettersen *et al.*, 2004), with the selected projection of resulting dimer model matching the observed 2D class averages. For 3D classification and refinement, a previously characterized structure of *S. cerevisiae* CMG (EMD-6535) was used as a starting model (Yuan *et al.*, 2016).

#### Author contributions

L.L. and Y.Z. performed most of the experiments except for the single-particle EM in Figures 6 and 7, which was carried out by J.Z. All the mass spectrometry analysis was performed by J-H.W., M-Q.D. and Z.L.

## Acknowledgments

We thank Dr. Costa for sharing unpublished data and discussion, Drs. Stephen Bell, and Li-Lin Du for reagents; Drs. Ning Gao, Hao Wu, Qun He, Yisui Xia and members of the Lou lab for helpful discussion and comments on the manuscript.

This work was supported by the National Natural Science Foundation of China 31630005, 31770084, 31771382 and 31271331; the National Basic Research Program (973 Program) of China (2014CB849801); Chinese Universities Scientific Fund 2015TC039 and 2014JD075; Opening Project of the State Key Laboratory of Microbial Resources; Program for Extramural Scientists of the State Key Laboratory of Agrobiotechnology 2018SKLAB6-5.

---

**Table S1.**
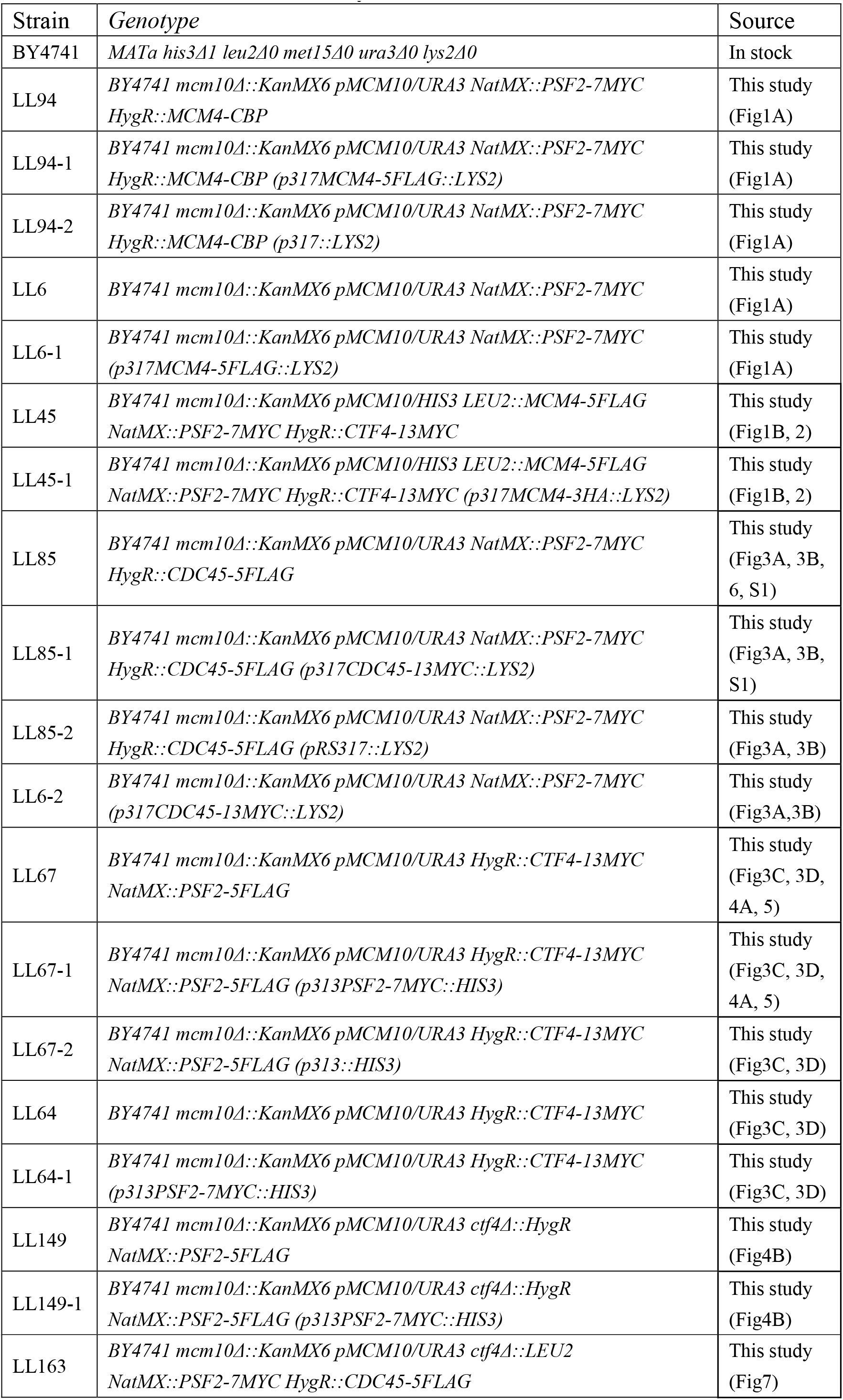
Strains used in this study.

**Table S2.** Plasmids used in this study

| Plasmid | Base plasmid/Genotype | Source |
| --- | --- | --- |
| pRS316-MCM10 | *ampr/URA3 MCM10* | This study |
| pRS317-MCM4-3HA | *ampr/LYS2 MCM4-3HA* | This study |
| pRS317-MCM4-5FLAG | *ampr/LYS2 MCM4-5FLAG* | This study |
| pRS317-CDC45-13MYC | *ampr/LYS CDC45-13MYC* | This study |
| pRS313-PSF2-7MYC | *ampr/HIS3 PSF2-7MYC* | This study |
| pRS313-MCM10 | *ampr/HIS3 MCM10* | This study |

## References

Abid Ali F, Renault L, Gannon J, Gahlon HL, Kotecha A, Zhou JC, Rueda D, Costa A (2016) Cryo-EM structures of the eukaryotic replicative helicase bound to a translocation substrate. Nature communications 7: 10708

Bell SP, Labib K (2016) Chromosome Duplication in Saccharomyces cerevisiae. Genetics 203: 1027–1067

Bleichert F, Botchan MR, Berger JM (2017) Mechanisms for initiating cellular DNA replication. Science 355: eaah6317

Bruck I, Kaplan DL (2015) The Replication Initiation Protein Sld3/Treslin Orchestrates the Assembly of the Replication Fork Helicase during S Phase. The Journal of biological chemistry 290: 27414–27424

Burgers PMJ, Kunkel TA (2017) Eukaryotic DNA Replication Fork. Annu Rev Biochem 86: 417–438

Chang YP, Wang G, Bermudez V, Hurwitz J, Chen XS (2007) Crystal structure of the GINS complex and functional insights into its role in DNA replication. Proc Natl Acad Sci USA 104: 12685–12690

Chi H, He K, Yang B, Chen Z, Sun RX, Fan SB, Zhang K, Liu C, Yuan ZF, Wang QH, Liu SQ, Dong MQ, He SM (2015) pFind-Alioth: A novel unrestricted database search algorithm to improve the interpretation of high-resolution MS/MS data. J Proteomics 125: 89–97

Costa A, Ilves I, Tamberg N, Petojevic T, Nogales E, Botchan MR, Berger JM (2011) The structural basis for MCM2-7 helicase activation by GINS and Cdc45. Nat Struct Mol Biol 18: 471–477

Costa A, Renault L, Swuec P, Petojevic T, Pesavento JJ, Ilves I, MacLellan-Gibson K, Fleck RA, Botchan MR, Berger JM (2014) DNA binding polarity, dimerization, and ATPase ring remodeling in the CMG helicase of the eukaryotic replisome. Elife 3: e03273

Coster G, Diffley JFX (2017) Bidirectional eukaryotic DNA replication is established by quasi-symmetrical helicase loading (vol 357, pg 314, 2017). Science 357: 314–318

Deegan TD, Yeeles JT, Diffley JF (2016) Phosphopeptide binding by Sld3 links Dbf4-dependent kinase to MCM replicative helicase activation. The EMBO journal 35: 961–973

Douglas ME, Ali FA, Costa A, Diffley JFX (2018) The mechanism of eukaryotic CMG helicase activation. Nature 555: 265–268

Douglas ME, Diffley JF (2016) Recruitment of Mcm10 to Sites of Replication Initiation Requires Direct Binding to the Minichromosome Maintenance (MCM) Complex. The Journal of biological chemistry 291: 5879–5888

Dua R, Edwards S, Levy DL, Campbell JL (2000) Subunit interactions within the Saccharomyces cerevisiae DNA polymerase epsilon (pol epsilon) complex. Demonstration of a dimeric pol epsilon. The Journal of biological chemistry 275: 28816–28825

Evrin C, Clarke P, Zech J, Lurz R, Sun J, Uhle S, Li H, Stillman B, Speck C (2009) A double-hexameric MCM2-7 complex is loaded onto origin DNA during licensing of eukaryotic DNA replication. Proc Natl Acad Sci USA 106: 20240–20245

Fang D, Cao Q, Lou H (2016) Sld3-MCM Interaction Facilitated by Dbf4-Dependent Kinase Defines an Essential Step in Eukaryotic DNA Replication Initiation. Front Microbiol 7: 885

Gambus A, Jones RC, Sanchez-Diaz A, Kanemaki M, van Deursen F, Edmondson RD, Labib K (2006) GINS maintains association of Cdc45 with MCM in replisome progression complexes at eukaryotic DNA replication forks. Nature cell biology 8: 358–366

Gambus A, van Deursen F, Polychronopoulos D, Foltman M, Jones RC, Edmondson RD, Calzada A, Labib K (2009) A key role for Ctf4 in coupling the MCM2-7 helicase to DNA polymerase alpha within the eukaryotic replisome. The EMBO journal 28: 2992–3004

Georgescu R, Yuan Z, Bai L, de Luna Almeida Santos R, Sun J, Zhang D, Yurieva O, Li H, O’Donnell ME (2017) Structure of eukaryotic CMG helicase at a replication fork and implications to replisome architecture and origin initiation. Proc Natl Acad Sci USA 114: E697–E706

Heller RC, Kang S, Lam WM, Chen S, Chan CS, Bell SP (2011) Eukaryotic origin-dependent DNA replication in vitro reveals sequential action of DDK and S-CDK kinases. Cell 146: 80–91

Ilves I, Petojevic T, Pesavento JJ, Botchan MR (2010) Activation of the MCM2-7 helicase by association with Cdc45 and GINS proteins. Molecular cell 37: 247–258

Itou H, Shirakihara Y, Araki H (2015) The quaternary structure of the eukaryotic DNA replication proteins Sld7 and Sld3. Acta Crystallogr D Biol Crystallogr 71: 1649–56

Kamada K, Kubota Y, Arata T, Shindo Y, Hanaoka F (2007) Structure of the human GINS complex and its assembly and functional interface in replication initiation. Nat Struct Mol Biol 14: 388–396

Li N, Zhai Y, Zhang Y, Li W, Yang M, Lei J, Tye B, Gao N (2015) Structure of the eukaryotic MCM complex at 3.8 A. Nature 524: 186–191

Miyazawa-Onami M, Araki H, Tanaka S (2017) Pre-initiation complex assembly functions as a molecular switch that splits the Mcm2-7 double hexamer. EMBO Reports 18: 1752–1761

Moyer SE, Lewis PW, Botchan MR (2006) Isolation of the Cdc45/Mcm2-7/GINS (CMG) complex, a candidate for the eukaryotic DNA replication fork helicase. Proc Natl Acad Sci USA 103: 10236–10241

Muramatsu S, Hirai K, Tak YS, Kamimura Y, Araki H (2010) CDK-dependent complex formation between replication proteins Dpb11, Sld2, Pol (epsilon}, and GINS in budding yeast. Genes & development 24: 602–612

O’Donnell ME, Li H (2018) The ring-shaped hexameric helicases that function at DNA replication forks. Nat Struct Mol Biol 25: 122–130

Pacek M, Tutter AV, Kubota Y, Takisawa H, Walter JC (2006) Localization of MCM2-7, Cdc45, and GINS to the site of DNA unwinding during eukaryotic DNA replication. Molecular cell 21: 581–587

Parker MW, Botchan MR, Berger JM (2017) Mechanisms and regulation of DNA replication initiation in eukaryotes. Crit Rev Biochem Mol Biol 52: 107–144

Perera RL, Torella R, Klinge S, Kilkenny ML, Maman JD, Pellegrini L (2013) Mechanism for priming DNA synthesis by yeast DNA polymerase alpha. Elife 2: e00482

Pettersen EF, Goddard TD, Huang CC, Couch GS, Greenblatt DM, Meng EC, Ferrin TE (2004) UCSF Chimera--a visualization system for exploratory research and analysis. J Comput Chem 25: 1605–1612

Quan Y, Xia Y, Liu L, Cui J, Li Z, Cao Q, Chen XS, Campbell JL, Lou H (2015) Cell-Cycle-Regulated Interaction between Mcm10 and Double Hexameric Mcm2-7 Is Required for Helicase Splitting and Activation during S Phase. Cell reports 13: 2576–2586

Remus D, Beuron F, Tolun G, Griffith JD, Morris EP, Diffley JF (2009) Concerted loading of Mcm2-7 double hexamers around DNA during DNA replication origin licensing. Cell 139: 719–730

Riera A, Barbon M, Noguchi Y, Reuter LM, Schneider S, Speck C (2017) From structure to mechanism-understanding initiation of DNA replication. Genes & development 31: 1073–1088

Scheres SH (2012) RELION: implementation of a Bayesian approach to cryo-EM structure determination. J Struct Biol 180: 519–530

Sengupta S, van Deursen F, de Piccoli G, Labib K (2013) Dpb2 integrates the leading-strand DNA polymerase into the eukaryotic replisome. Current biology : CB 23: 543–552

Sheu YJ, Stillman B (2006) Cdc7-Dbf4 phosphorylates MCM proteins via a docking site-mediated mechanism to promote S phase progression. Molecular cell 24: 101–113

Sheu YJ, Stillman B (2010) The Dbf4-Cdc7 kinase promotes S phase by alleviating an inhibitory activity in Mcm4. Nature 463: 113–117

Siddiqui K, On KF, Diffley JF (2013) Regulating DNA replication in eukarya. Cold Spring Harb Perspect Biol 5: a012930

Simon AC, Zhou JC, Perera RL, van Deursen F, Evrin C, Ivanova ME, Kilkenny ML, Renault L, Kjaer S, Matak-Vinkovic D, Labib K, Costa A, Pellegrini L (2014) A Ctf4 trimer couples the CMG helicase to DNA polymerase alpha in the eukaryotic replisome. Nature 510: 293–297

Sun J, Shi Y, Georgescu RE, Yuan Z, Chait BT, Li H, O’Donnell ME (2015) The architecture of a eukaryotic replisome. Nat Struct Mol Biol 22: 976–982

Sun J, Yuan Z, Georgescu R, Li H, O’Donnell M (2016) The eukaryotic CMG helicase pumpjack and integration into the replisome. Nucleus 7: 146–154

Swuec P, Costa A (2017) DNA replication and inter-strand crosslink repair: Symmetric activation of dimeric nanomachines? Biophys Chem 225: 15–19

Tanaka S, Araki H (2013) Helicase activation and establishment of replication forks at chromosomal origins of replication. Cold Spring Harb Perspect Biol 5: a010371

Tang G, Peng L, Baldwin PR, Mann DS, Jiang W, Rees I, Ludtke SJ (2007) EMAN2: an extensible image processing suite for electron microscopy. J Struct Biol 157: 38–46

Villa F, Simon AC, Ortiz Bazan MA, Kilkenny ML, Wirthensohn D, Wightman M, Matak-Vinkovic D, Pellegrini L, Labib K (2016) Ctf4 Is a Hub in the Eukaryotic Replisome that Links Multiple CIP-Box Proteins to the CMG Helicase. Molecular cell 63: 385–396

Watson JD, Crick FH (1953) Molecular structure of nucleic acids; a structure for deoxyribose nucleic acid. Nature 171: 737–738

Xia Y, Niu Y, Cui J, Fu Y, Chen XS, Lou H, Cao Q (2015) The Helicase Activity of Hyperthermophilic Archaeal MCM is Enhanced at High Temperatures by Lysine Methylation. Front Microbiol 6: 1247

Xu Y, Gristwood T, Hodgson B, Trinidad JC, Albers SV, Bell SD (2016) Archaeal orthologs of Cdc45 and GINS form a stable complex that stimulates the helicase activity of MCM. Proc Natl Acad Sci USA 113: 13390–13395

Yardimci H, Loveland AB, Habuchi S, van Oijen AM, Walter JC (2010) Uncoupling of sister replisomes during eukaryotic DNA replication. Molecular cell 40: 834–840

Yeeles JT, Deegan TD, Janska A, Early A, Diffley JF (2015) Regulated eukaryotic DNA replication origin firing with purified proteins. Nature 519: 431–435

Yuan Z, Bai L, Sun J, Georgescu R, Liu J, O’Donnell ME, Li H (2016) Structure of the eukaryotic replicative CMG helicase suggests a pumpjack motion for translocation. Nat Struct Mol Biol 23: 217–224

Zhou JC, Janska A, Goswami P, Renault L, Abid Ali F, Kotecha A, Diffley JFX, Costa A (2017) CMG-Pol epsilon dynamics suggests a mechanism for the establishment of leading-strand synthesis in the eukaryotic replisome. Proc Natl Acad Sci USA 114: 4141–4146

